# The Non-Human Primate Sample Size Calculator: a design tool for longitudinal research projects

**DOI:** 10.64898/2026.08.06.743285

**Authors:** Hannah C. Ainsworth, Hillary F. Huber, Ellen E. Quillen, Brooke Justice, Patricia L. Warren, Timothy D. Howard, Laura A. Cox

**Affiliations:** Department of Biostatistics and Data Science, Wake Forest University School of Medicine, Winston-Salem, North Carolina, USA; Center for Precision Medicine, Wake Forest University School of Medicine, Winston-Salem, North Carolina, USA; Southwest National Primate Research Center, Texas Biomedical Research Institute, San Antonio, Texas, USA; Section of Molecular Medicine, Department of Internal Medicine, Wake Forest University School of Medicine, Winston-Salem, North Carolina, USA; Section on Comparative Medicine, Department of Pathology, Wake Forest University School of Medicine, Winston-Salem, North Carolina, USA; Department of Biochemistry, Wake Forest University School of Medicine, Winston-Salem, North Carolina, USA; Section on Molecular Medicine, Department of Internal Medicine, Wake Forest University School of Medicine, Winston-Salem, North Carolina, USA

## Abstract

Nonhuman primates (NHP) are crucial models of human health and disease and offer the opportunity to bridge the gap between basic research and the clinic. Given the ethical considerations, high costs, and logistical constraints associated with NHP studies, careful planning is essential to maximize scientific rigor while minimizing animal use. Statistical power calculations, to determine optimal sample size, require knowing the expected number of animals at each timepoint, a challenge for longitudinal studies that must account for natural deaths or disease during a study. Here, we leveraged data from a recent study detailing NHP life- and healthspans to develop a web-based NHP study design tool, available at midas.wakehealth.edu. For 11 NHP species relevant to biomedical research, users can provide study details to generate estimates of sample counts based on natural healthspan trajectories. We envision this tool being used by colony managers and investigators for both study design and monitoring colony health over the course of a study.

## INTRODUCTION

Nonhuman primates (NHP) are crucial models of human health and disease because they share profound genetic, physiological, and behavioral similarities with humans, more so than any other animal model, making them uniquely suited to bridge the gap between basic research and the clinic. Their likeness enables scientists to study complex biological systems, diseases, and therapeutics in ways that are directly translatable to humans, supporting major breakthroughs in areas such as neuroscience, immunology, infectious disease, and reproductive health^1–9^. Research with NHP has been central to the development of vaccines, treatments for neurodegenerative diseases, cancer therapies, and reproductive medicine, vastly improving medical outcomes for people around the world^1–9^.

Given the ethical considerations, high costs, and logistical constraints associated with NHP studies, animal availability is often limited and thus careful planning is essential to maximize scientific rigor while minimizing animal use^10^. During study design, statistical power calculations are essential for balancing minimum sample size with the ability to detect statistical associations^11^. This analysis aids researchers in designing experiments that are adequately powered to detect meaningful effects, ensure reproducibility, and uphold ethical standards by preventing unnecessary use of these valuable animals^12,13^. However, proper power analysis for a study requires knowledge of expected number of animals at each timepoint. This estimate can be particularly challenging for longitudinal studies that must account for the number of animals that might be lost to natural death or disease during a study. This information is critical for determining the starting sample size and for monitoring whether study subject drop-out is at an expected level.

Leveraging data from a recent study detailing survivorship across multiple NHP species^14^, we developed a web-based sample calculator, available at **midas.wakehealth.edu** (see ‘availability statement’). This tool allows users to choose a species (out of 11 NHP choices), age at study start, sex of study subjects, duration of the study, and number of animals necessary at either the start or end of the study. We envision this tool being used by colony managers and investigators for both study design and monitoring colony health over the course of a study.

## DESCRIPTION

The source dataset was assembled from our previously published study on the lifespans and healthspans of research NHP from 15 NHP research institutions in the United States and Indonesia.^14^ Within the sample calculator, 11 species are included (**Table 1**) for which sufficient sample size was available for prediction, and for which permission was obtained for including the data. Sample processing was previously described^14^. Briefly, animal records were filtered to animals that survived to at least adulthood, as defined by the National Institutes of Health Nonhuman Primate Evaluation and Analysis table of NHP life stages^15^, as presented in **Table 1**. Institutional codes were reviewed to retain only animals that died of natural causes or were euthanized for clinical/health reasons. Because analyses did not include living animals, a species-specific date of birth threshold was implemented to avoid bias from earlier deaths among recently-born animals (described in detail^14^). All data are stored within MIDAS (Monkey Inventory and Data Management of Samples). MIDAS is a multi-NHP species resource within the LabKey Laboratory Information Management System (LIMS).

**Table 1.** Species and sample sizes on which the survivorship estimator tool is based.

| **Common Name** | **Species name** | **Age at Adulthood\*** | **Male** | **Female** |
| --- | --- | --- | --- | --- |
| Baboon | *Papio hamadryas* spp. | 4 years | 334 | 669 |
| Bonnet macaque | *Macaca radiata* | 4 years | 19 | 43 |
| Common marmoset | *Callithrix jacchus* | 1.5 years | 378 | 453 |
| Coppery titi monkey | *Plecturocebus cupreus* | 4 years | 32 | 33 |
| Cotton-top tamarin | *Saguinus oedipus* | 2.5 years | 155 | 191 |
| Cynomolgus macaque | *Macaca fascicularis* | 4 years | 82 | 132 |
| Japanese macaque | *Macaca fuscata* | 4 years | 174 | 196 |
| Pig-tailed macaque | *Macaca nemestrina* | 4 years | 173 | 596 |
| Rhesus macaque | *Macaca mulatta* | 4 years | 2465 | 5742 |
| Squirrel monkey | *Saimiri* spp. | 4 years | 53 | 47 |
| Vervet/African green | *Chlorocebus aethiops sabaeus* | 4 years | 60 | 144 |

\* Age at adulthood based on National Institutes of Health Nonhuman Primate Evaluation and Analysis table of NHP life stages<sup>15</sup>

For each species, by sex, Kaplan-Meier survivorship estimates were previously computed as^14,16^:

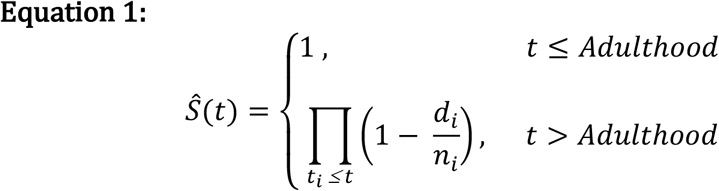

with *t*_*i*_ as time when at least one event happened, *d*_*i*_ the number of deaths that occurred until time *t*_*i*_ and, *n*_*i*_ the number of animals known to have survived to this time point. As described above, adulthood was defined on a species-specific basis via NIH guidelines^14^.

With these survivorship estimates, the number of animals for a particular study design can be computed with user-specified parameters. Recalling that all survival estimates are based on adult (NHP-specific) data, computing the number of animals at the end (**t**_**end**_) of a multi-year study that starts at adulthood can be achieved by Equation 2:

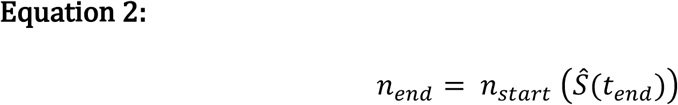

Alternatively, computing the number of animals at the end (t_end_) of a multi-year study, which begins after adulthood (**t**_**start**_) requires accounting for the conditional probability of assumed survival to **t**_**start**_. That is, an animal that has already survived to an age beyond adulthood would have an increased probability to surviving to **t**_**end**_. This is computed as shown in Equation 3:

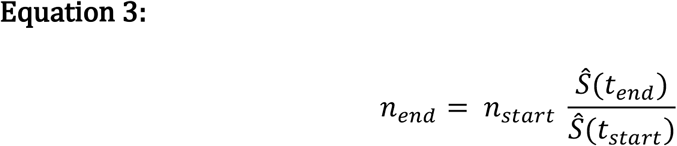

These calculations are handled by a graphical user interface (**Figure 1**), which is freely available via the “NHP Study Design Tool” links at **midas.wakehealth.edu**. For access, users are requested to provide basic information including name, affiliation, and email address to help track the usage and utility of this resource. The NHP Sample Calculator was built using a custom JavaScript which accesses R-4.5.3 to use ggplot for visualization of the Kaplan-Meier survival curves.

**Figure 1:**
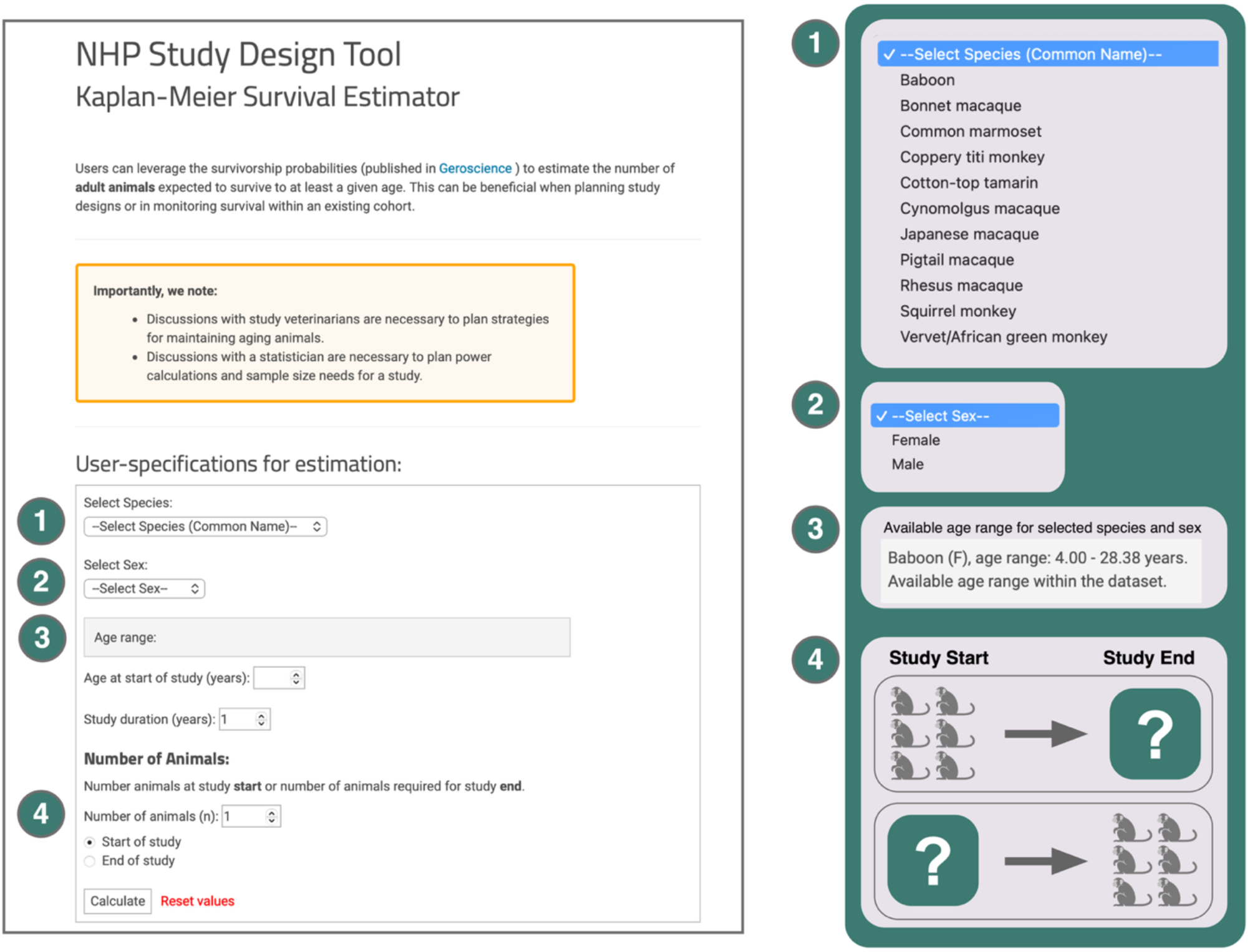
Overview of the NHP Sample Calculator. A screenshot of the NHP Study Design Tool available on MIDAS. Menu options for species and sex are shown. Upon selection of species and sex, the calculator will automatically update with the available age range in the sourced dataset. After selecting age at start of study, duration of study, and number of animals, the user selects whether the provided number of animals corresponds to the study end or study start.

## EXAMPLE

We describe the two primary use case scenarios for the NHP Sample Calculator. Use Case 1 focuses on computing sample estimates for the end of a study and Use Case 2 focuses on computing the number of animals needed at the start of a study. Both follow the same user experience but provide distinct pieces of information. Each use case is presented with an example scenario.

### Use Case 1: Determine estimated animal survivorship over a period of time

This use case has implications for an investigator who may want to compare how survivorship within their study compares to natural and health-related survivorship in a matched species. This may also be informative for a colony manager who needs to plan how many animals are expected to survive to a particular age. For example (**Figure 2**), a researcher has been following a group of marmosets since 3 years of age (“Age at start of study”). The study started with 16 male marmosets. The group has reached 6 years old (“Study duration (years)” = 3) and there are 10 marmosets remaining. How does this survival compare to survivorship estimates provided by the NHP lifespan study?

**Figure 2:**
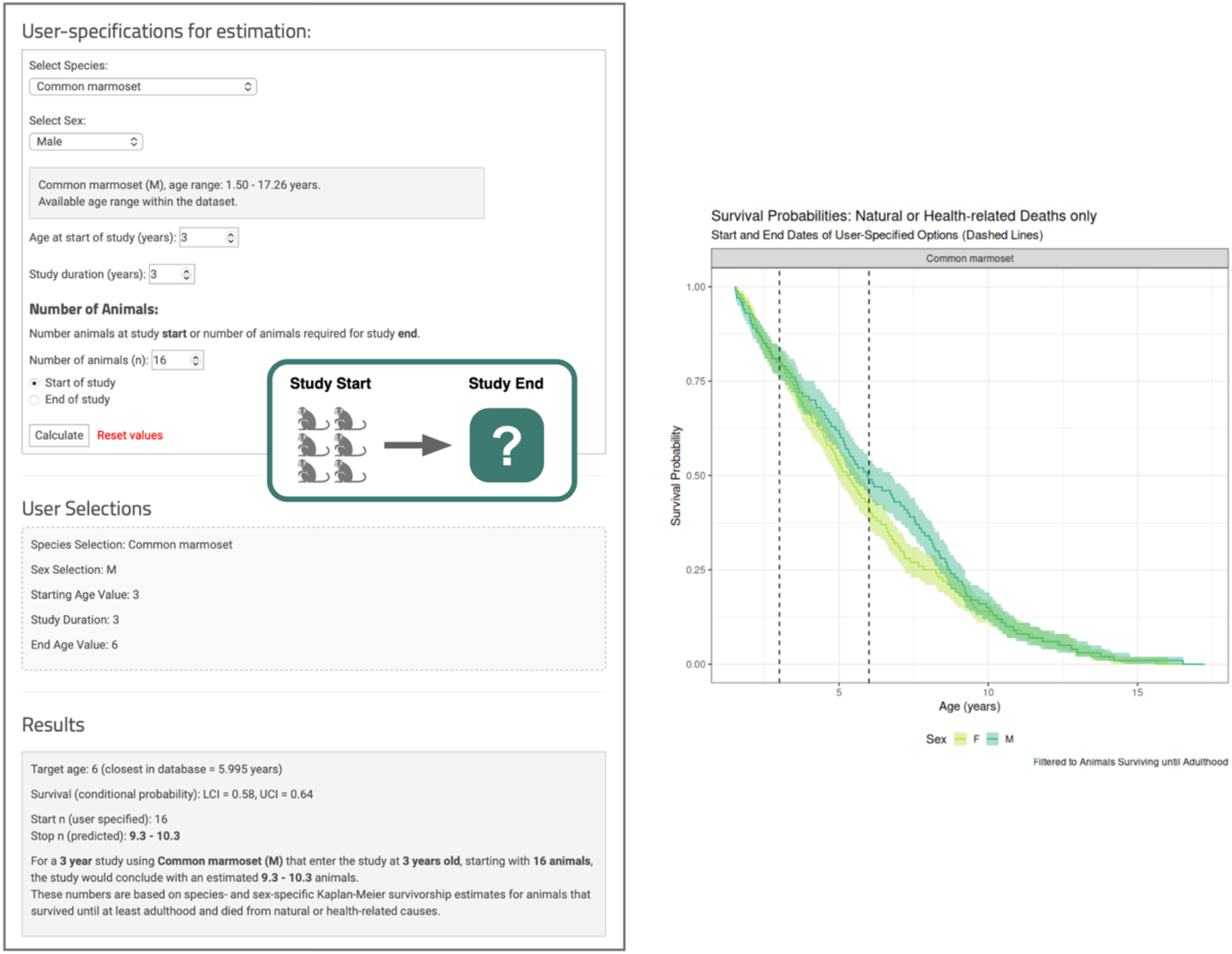
Use Case, Example 1: Estimating the number of animals at the end of a study. Calculator workflow for options described in Example 1. Upon selection of all options, a summary of the user-input is provided along with a results statement. The Kaplan-Meier curves for the selected species are also provided, with the start and end ages of NHP in the study denoted as vertical dashed lines. Both sexes are provided in plot to further help inform study design considerations. Here, calculations are based on the conditional probability of animals surviving to at least 3 years of age, which is after onset of adulthood (1.5 years).

In selecting the appropriate parameters (species, sex, and time span of the study), the NHP Sample Calculator shows that the expected number of male marmosets, based on a 95% confidence, would be 9.3-10.3 animals, thus supporting that the current survival of 10 marmosets aligns with existing data. This workflow would also support statistical study design considerations. If a longitudinal study is limited to using a certain number of animals, this tool could help inform how many animals are expected to survive to the end of the study. This would also contribute to statistical power calculations for detectable effect sizes.

### Use Case 2: Determine number of animals needed at the start of a longitudinal study

This use case has critical implications for study design and statistical power calculations. For instance, working with a statistician, an investigator may learn they need to have 6 animals per group to achieve sufficient statistical power at the end of their 4-year study (**Figure 3**). If using female cynomolgus macaques that will enter the study at adulthood (4 years old), how many animals should begin the study? Akin to the previous example, the user would select the appropriate parameters (species, sex, and time span of the study) but instead select “at end of study.” In doing so, the NHP Sample Calculator indicates that if ending a 4-year study with 6 female cynomolgus macaques is necessary, the researcher should plan to start with 9.4 -12.8 animals, which spans the 95% confidence interval.

**Figure 3:**
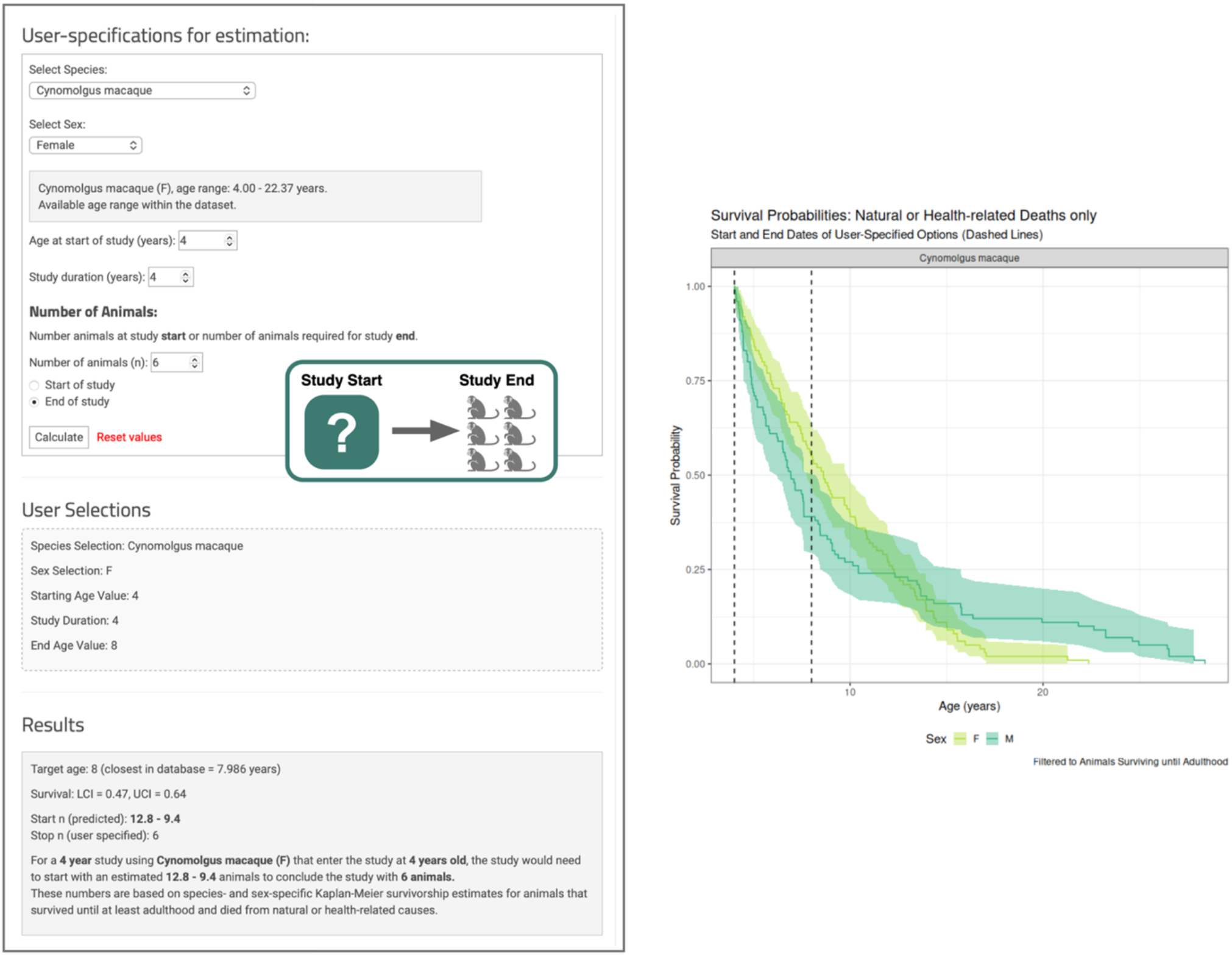
Use Case, Example 2: Estimating the number of animals needed at the beginning of a study, given a target sample size at study’s end. Calculator workflow for options described in Example 2. Upon selection of all options, a summary of the user-input is provided along with a results statement. The Kaplan-Meier curves for the selected species are also provided, with the start and end ages of NHP in the study denoted as vertical dashed lines. Both sexes are provided in the plot to further inform study design considerations. In this example, the starting age occurs at adulthood, with a survivorship probability of 1.

## COMPARISON AND CRITIQUE

Estimation of NHP survivorship is essential for study design, colony management, and data interpretation. Yet, existing methods rely largely on estimates based on a single published study^14,17,18^, or even a single published animal^19^. Complicating the issue, many NHP lifespan resources give only the maximum known lifespan for a species, rather than the average or expected lifespan^19–22^, akin to reporting human lifespan as 120 years rather than somewhere in the seventh or eighth decade of life (population dependent). Some NHP study subjects will inevitably die of natural causes or require humane euthanasia for causes unrelated to the studies in which they participate (e.g., cancer, infectious disease), affecting study sample size and data interpretation. Our NHP sample calculator is the first web-based tool to provide adult survivorship estimates across multiple NHP species, based on decades of data, across multiple NHP research centers. Our tool provides a user-friendly resource to automate sample size estimation from previously published Kaplan-Meier survivorship estimates.

There are several noted limitations to this tool. First, the data are sourced from adult animals and thus not informative for studies utilizing juvenile animals. A future goal of the project is to assemble and analyze juvenile data. Second, all analyses were limited to animals that died of natural or health-related reasons. Although survivorship probabilities can be computed with censored data (here, meaning a death due to non-health related reasons, *right censored*), in our previous analyses of these data^14^, we found censoring to be informative (e.g., confounded with sex) and not appropriate for statistical inference. Consequently, usage of only natural or health-related deaths yielded small sample sizes for some species (e.g., bonnet macaque or coppery titi monkey), which affects the confidence of the sample size estimate about the point estimate; thus, all results include 95% confidence intervals. Third, this tool is likely only relevant for animals in biomedical research conditions. It is not applicable to wild NHPs, which do not receive ongoing medical care. It may also be imprecise for zoo or sanctuary NHPs, who live under different conditions than research NHPs. Finally, individual studies may have factors that affect survivorship beyond natural causes; such factors would not be reflected by the calculator. As such, we caution that 1) discussions with study veterinarians are necessary to plan strategies for maintaining aging animals, and 2) discussions with a statistician are necessary to plan power calculations and sample size needs for a study.

We expect this tool to be valuable for NHP study design and interpretation. It can directly contribute to the “three Rs” of animal research to replace, reduce, and refine the use of animals by enabling researchers to more accurately predict participant survival rates and thus optimize experimental design. By refining estimates for necessary group sizes based on likely outcomes, such a tool helps “reduce” the total number of animals used, preventing both underpowered and excessively large studies. Better survivorship predictions also enable “refinement” of experiments by avoiding endpoint scenarios where too few survivors lead to repeat or failed studies, in turn sparing animals unnecessary procedures or distress. Although not a direct replacement strategy, improved predictions from a survivorship tool support the “replacement” principle by minimizing the need for additional cohorts and facilitating more effective use of non-animal alternatives when feasible. In sum, a survivorship estimator embodies the three Rs by ensuring scientific rigor while upholding high ethical standards in animal research.

## AVAILABILITY

The sample calculator can be found via the “NHP Study Design Tool” links available at midas.wakehealth.edu. For access, users are requested to provide basic information including name, affiliation, and email address to help track the usage and utility of this resource.

## ACKNOWLEDGMENTS

We thank the beta testers who provided feedback on use of the calculator during development. This work was supported by the National Institute on Aging: R33 AG073733 (LAC) and R01AG087957 (EEQ). Initial testing of code and computations were performed using the Wake Forest University (WFU) High Performance Computing Facility, a centrally managed computational resource available to WFU researchers including faculty, staff, students, and collaborators.

## REFERENCES

1. Friedman H, Ator N, Haigwood N, Newsome W, Allan JS, Golos TG, Kordower JH, Shade RE, Goldberg ME, Bailey MR, Bianchi P. The Critical Role of Nonhuman Primates in Medical Research. Pathog Immun. 2017;2(3):352–365. doi:10.20411/pai.v2i3.186

2. Phillips KA, Bales KL, Capitanio JP, Conley A, Czoty PW, ‘t Hart BA, Hopkins WD, Hu SL, Miller LA, Nader MA, Nathanielsz PW, Rogers J, Shively CA, Voytko ML. Why primate models matter. Am J Primatol. 2014;76(9):801–827. doi:10.1002/ajp.22281

3. Nakamura T, Fujiwara K, Saitou M, Tsukiyama T. Non-human primates as a model for human development. Stem Cell Rep. 2021;16(5):1093–1103. doi:10.1016/j.stemcr.2021.03.021

4. Choudhury GR, Kim J, Frost PA, Bastarrachea RA, Daadi MM. Nonhuman primate model in clinical modeling of diseases for stem cell therapy. Brain Circ. 2016;2(3):141. doi:10.4103/2394-8108.192524

5. Simerly CR, Castro CA, Jacoby E, Grund K, Turpin J, McFarland D, Champagne J, Jimenez JB, Frost P, Bauer C, Hewitson L, Schatten G. Assisted Reproductive Technologies (ART) with Baboons Generate Live Offspring: A Nonhuman Primate Model for ART and Reproductive Sciences. Reprod Sci. 2010;17(10):917–930. doi:10.1177/1933719110374114

6. Vallender EJ, Miller GM. Nonhuman Primate Models in the Genomic Era: A Paradigm Shift. ILAR J. 2013;54(2):154–165. doi:10.1093/ilar/ilt044

7. Vallender EJ, Hotchkiss CE, Lewis AD, Rogers J, Stern JA, Peterson SM, Ferguson B, Sayers K. Nonhuman primate genetic models for the study of rare diseases. Orphanet J Rare Dis. 2023;18:20. doi:10.1186/s13023-023-02619-3

8. Petroff RL, Grant KS, Burbacher TM. The Role of Nonhuman Primates in Neurotoxicology Research: Preclinical Models and Experimental Methods. Curr Protoc. 2023;3(3):e698. doi:10.1002/cpz1.698

9. Miller LA, Royer CM, Pinkerton KE, Schelegle ES. Nonhuman Primate Models of Respiratory Disease: Past, Present, and Future. ILAR J. 2017;58(2):269–280. doi:10.1093/ilar/ilx030

10. Huber HF, Jenkins SL, Li C, Nathanielsz PW. Strength of nonhuman primate studies of developmental programming: review of sample sizes, challenges, and steps for future work. J Dev Orig Health Dis. 2020;11(3):297–306. doi:10.1017/S2040174419000539

11. Casella G, Berger R. Statistical Inference. 2nd ed. Chapman and Hall/CRC; 2024. doi:10.1201/9781003456285

12. Cauvin AJ, Peters C, Brennan F. Advantages and Limitations of Commonly Used Nonhuman Primate Species in Research and Development of Biopharmaceuticals. Nonhum Primate Nonclinical Drug Dev Saf Assess. Published online 2015:379–395. doi:10.1016/B978-0-12-417144-2.00019-6

13. Bliss-Moreau E, Amara RR, Buffalo EA, Colman RJ, Embers ME, Morrison JH, Quillen EE, Sacha JB, Roberts CT, National Primate Research Center Consortium Rigor and Reproducibility Working Group. Improving rigor and reproducibility in nonhuman primate research. Am J Primatol. 2021;83(12):e23331. doi:10.1002/ajp.23331

14. Huber HF*, Ainsworth HC*, Quillen EE, Salmon A, Corinna Ross, Azhar AD, Bales K, Basso MA, Coleman K, Colman R, Darusman HS, Hopkins W, Hotchkiss CE, Jorgensen MJ, Kavanagh K, Li C, Mattison JA, Nathanielsz PW, Saputro S, Scorpio DG, Sosa PM, Vallender EJ, Wang Y, Zeiss CJ, Shively CA, Cox LA. Comparative lifespan and healthspan of nonhuman primate species common to biomedical research. GeroScience. 2025;47:135–151. doi:10.1007/s11357-024-01421-8

15. Feister AJ, DiPietrantonio A, Yuenger J, Ireland K, Rao A. Nonhuman Primate Evaluation and Analysis Part 1: Analysis of Future Demand and Supply | Office of Research Infrastructure Programs (ORIP) – DPCPSI – NIH. 2018. Accessed August 9, 2023. https://orip.nih.gov/nonhuman-primate-evaluation-and-analysis-part-1-analysis-future-demand-and-supply

16. Hosmer, Jr. DW, Lemeshow S, May S. Applied Survival Analysis: Regression Modeling of Time-to-Event Data. John Wiley & Sons; 2008.

17. Hadley EC, Nguyen CV, Gerald MS, Mattison JA, Moro M, Simmons JM, Alberts SC, Austad SN, Babbitt CC, Barton RA, Baudisch A, Cohen AA, de Magalhães JP, Thompson ME, Ferrucci L, Finch CE, Franchini LF, Girke T, Gladyshev VN, Goyal M, Hild S, Lammey ML, Melin AD, Miller RA, Morrison JH, Pontzer H, Rogers J. Research on Determinants of Species Differences in Human and Nonhuman Primate Life Spans and Health Spans. Published online 2021. https://www.nia.nih.gov/sites/default/files/2023-02/report_determinants_of_species_differences.pdf

18. Riddle NC, Biga PR, Bronikowski AM, Walters JR, Wilkinson GS, Duan JE, Gamble A, Larschan E, Meisel RP, Singh R, Webb A, IISAGE Consortium. Comparative analysis of animal lifespan. GeroScience. 2024;46(1):171–181. doi:10.1007/s11357-023-00984-2

19. Weigl R. Longevity of Mammals in Captivity; from the Living Collections of the World. Kleine Senckenberg-Reihe; 2005. Accessed March 1, 2021. https://www.schweizerbart.de/publications/detail/isbn/9783510613793/Longevity-of-mammals-in-captivity-from-the-Living-Collections-of-the-world

20. Zimmermann E, Radespiel U. Primate Life Histories. In: Henke W, Tattersall I, eds. Handbook of Paleoanthropology. Springer; 2015:1527–1592. doi:10.1007/978-3-642-39979-4_38

21. Tacutu R, Thornton D, Johnson E, Budovsky A, Barardo D, Craig T, Diana E, Lehmann G, Toren D, Wang J, Fraifeld VE, de Magalhães JP. Human Ageing Genomic Resources: new and updated databases. Nucleic Acids Res. 2018;46(D1):D1083–D1090. doi:10.1093/nar/gkx1042

22. Hakeem AY, Sandoval GR, Jones M, Allman JM. Brain and life span in primates. In: Handbook of the Psychology of Aging. Academic Press, Inc; 1996:78–104.

